# Loss of daylength sensitivity by splice site mutation in Cannabis

**DOI:** 10.1101/2023.03.10.532103

**Authors:** Keegan M. Leckie, Jason Sawler, Paul Kapos, John O. Mackenzie, Ingrid Giles, Katherine Baynes, Jessica Lo, Jose M Celedon, Gregory J. Baute

## Abstract

Adaptations to high latitude photoperiods have been under positive selection during the domestication of many short-day (SD) flowering crops. Photoperiod insensitivity (auto-flowering) in drug-type *Cannabis sativa* circumvents the need for SD flowering requirements making outdoor cultivation in high latitudes possible. However, the benefits of photoperiod insensitivity are counterbalanced by low cannabinoid contents and poor flower quality in auto-flowering genotypes. Despite recent legalization in some countries, a mechanistic understanding of photoperiod insensitivity in cannabis is still lacking. Herein, we identify a splice site mutation within *PSEUDO-RESPONSE REGULATOR 37 (CsPRR37*) in auto-flowering cannabis that causes photoperiod insensitivity. Using a combination of GWAS, fine mapping, and gene expression analyses, our results strongly indicate *CsPRR37* as the most likely candidate for causing photoperiod insensitivity. Research into the pervasiveness of this mutation and others effecting flowering time will help elucidate its domestication history and advance cannabis breeding towards a more sustainable outdoor cultivation system.

## Introduction

The transition from vegetative growth to flowering is one of the most important developmental timepoints in a plant’s lifecycle. As changes in daylength correspond to changing seasons, many plants have evolved strong photoperiod sensitivity to determine when to flower. This photoperiod sensitivity can either be triggered by short days (SD) where flowering occurs when nights are sufficiently long or by long days (LD) in plants where daylength needs to exceed a critical threshold to induce flowering. Selective pressures including latitude and temperature, likely shaped photoperiod requirements (Ray and Alexander, 1966). Some plants in higher latitudes flower under LD in the summer to escape low temperatures in late spring or early fall and ensure enough time for seed maturation. At lower latitudes, some plants flower under SD in the spring to avoid the extreme heat of the summer (Samach and Coupland, 2000). Many domesticated crops have benefited from loss or change of photoperiod requirements that allow them to flower under day-length conditions different from their center of origin (see Nakamichi 2015). In fact, modification of flowering time is one of the most common domestication syndromes. The ancestral progenitors of many economically important crops including Sorghum, maize, potato, and *Japonica* rice are thought to have originated in lower latitudes, and through domestication acquired the ability to flower (or tuberization in potato) in LD facilitating their cultivation in new environments at higher latitudes (Fujino et al., 2022; Huang et al., 2018; Hardigan et al., 2017; Klein et al., 2015; Koo et al., 2013; Hung et al., 2012).

*Cannabis sativa* is an economically important SD crop likely originating in north-western China (Ren et al., 2021; McPartland et al., 2019). Several domesticated forms exist including those grown for seed and fiber, referred to as hemp, and those grown for medicinal and recreational purposes, referred to as drug-type cannabis. Drug-type cannabis can be defined by having high levels of total cannabinoids, regardless of their composition (CBD-dominant, THC-dominant, balanced CBD:THC, etc.). Cultivation of drug-type cannabis is typically done in indoor facilities using LD of 18 hours of light and 6 hours of dark (18h:6h) for vegetative growth followed by a switch to SD (12h:12h) to induce flowering. Interestingly, most drug-type cannabis will remain in a vegetative state indefinitely under LD as long as there are no other stresses that induce flowering. However, within both hemp and drug-type cannabis there are genotypes that flower after completing their juvenile phase regardless of day length. These day length neutral genotypes are typically referred to as “auto-flowering” cannabis. In addition to domesticated varieties, natural populations of cannabis from northern China have also been shown to flower under LD (Chen et al. 2022) indicating that the trait has an adaptive value and may be widespread in the wild.

The ability for plants to integrate daylength (and other environmental signals) in the onset of flowering is mediated by the circadian clock. The clock allows plants to anticipate seasonal changes by synchronizing development with the time of the year. In *Arabidopsis*, where the topic is most well studied, the circadian clock consists of interlocking transcription-translation feedback-loops that collectively compose a self-regulating system (Hsu and Harmer 2014). Clock genes exhibit oscillating expression patterns whose peak expression is localized to specific times of the day. This is most evident in the expression patterns of morning complex genes *CCA1/LHY* (Wang and Tobin, 1998; Schaffer et al., 1998) and the *TOC1* quintet of pseudo-response regulators (PRR) (Strayer et al., 2000; Matsushika et al., 2000). Other important genes involved in the circadian clock include evening complex (EC) genes *ELF3, ELF4*, and *LUX*, (Nusinow et al. 2011) and clock associated gene *GIGANTEA (GI*) (Fowler et al., 1999; Park et al., 1999), among others. The oscillation patterns of clock genes change in response to photoperiod, thereby regulating downstream floral induction machinery including circadian regulated *CONSTANS (CO*) and florigen *FLOWERING LOCUS T (FT*) (Suarez-Lopez et al., 2001; Yanovsky and Kay, 2002; Mizoguchi et al., 2005; Nakamichi et al., 2007). Disruption of proper gene expression of any one of these core clock components has been shown to alter flowering time (Nakamichi et al., 2007; Murakami et al., 2004; Yamamoto et al., 2003; Mizoguchi et al., 2002; Sato et al., 2002; Folwer et al., 1999; Somers et al., 1998).

With the increasing demand of hemp cultivation (Parvez et al., 2021) and ongoing changes in the legality of high THC drug-type cannabis, particularly in countries of high latitudes, flowering time has become a focus in cannabis breeding. Typical drug type cannabis requires 8 weeks of warm growing conditions to mature flower after the initiation of flowering. With photoperiod sensitive genotypes grown in high latitude regions the onset of flowering occurs in late summer, which has a high risk of encountering inclement weather before harvest. This is one of the reasons why the majority of drug-type cannabis grown in Canada is done indoors, or in greenhouses equipped with blackout curtains, a practise that is energy intensive and costly for growers, consumers and the environment. Short season, day length neutral cannabis allows growers to grow outdoors at whichever time of year best suites their bioregion.

The molecular mechanisms controlling flowering time in cannabis remains largely unexplored. Recent work done by Pan et al. (2021) observed that many of the *CO* homologues in cannabis change their expression profiles in LD compared to SD. Expression differences in *CO* were also observed between early and late flowering varieties. As *CO* is the prime regulator of *FT* in *Arabidopsis* and functions downstream of the circadian clock, these results suggest that altered clock outputs likely regulate flowering time in cannabis. Recently, Toth et al. (2022) mapped two cannabis QTLs involved in flowering time, one responsible for auto-flowering in cannabis which was mapped to a 5.2 Mb region of chromosome 1 (NC_044371.1, CBDRx reference genome). Consistent with Toth et al. (2022), we found the auto-flower trait to segregate as a single recessive locus in multiple F2 populations. Herein, we identify the putative causal gene of the auto-flowering QTL through a combination of gene expression and population genetics approaches. Our analyses indicate that a splice site mutation in *CsPRR37* causes auto-flowering in drug type cannabis genotypes. We show that auto-flowering plants have an altered circadian clock under LD which likely leads to flowering. Given the preponderance of flowering time alleles caused by aberrant PRR genes in domesticated crops (Koo et al., 2013; Pin et al., 2012; Murphy et al., 2011; Beales et al., 2007; Turner et al., 2005; Murakami et al., 2005), including SD species grown at high latitudes, a deeper understanding of the auto-flowering trait in cannabis may help to elucidate its domestication history.

## Results

### Auto-flowering is a Single Locus Recessive Allele in Linkage

To map the auto-flowering locus within the cannabis genome, Genome Wide Association (GWAS) was performed using a mixed linear model with both kinship and population structure accounted for. Genotype-by-Sequencing (GBS) data from a population of 231 photoperiod sensitive and 47 auto-flowering genotypes were used with the Purple Kush assembly as reference (van Bakel et a., 2011). SNPs associated with auto-flowering were identified spanning a large region on chromosome CM010796 (annotated as chromosome 5 in Purple Kush Assembly v.4) (Figure 1a).

**Figure 1.**
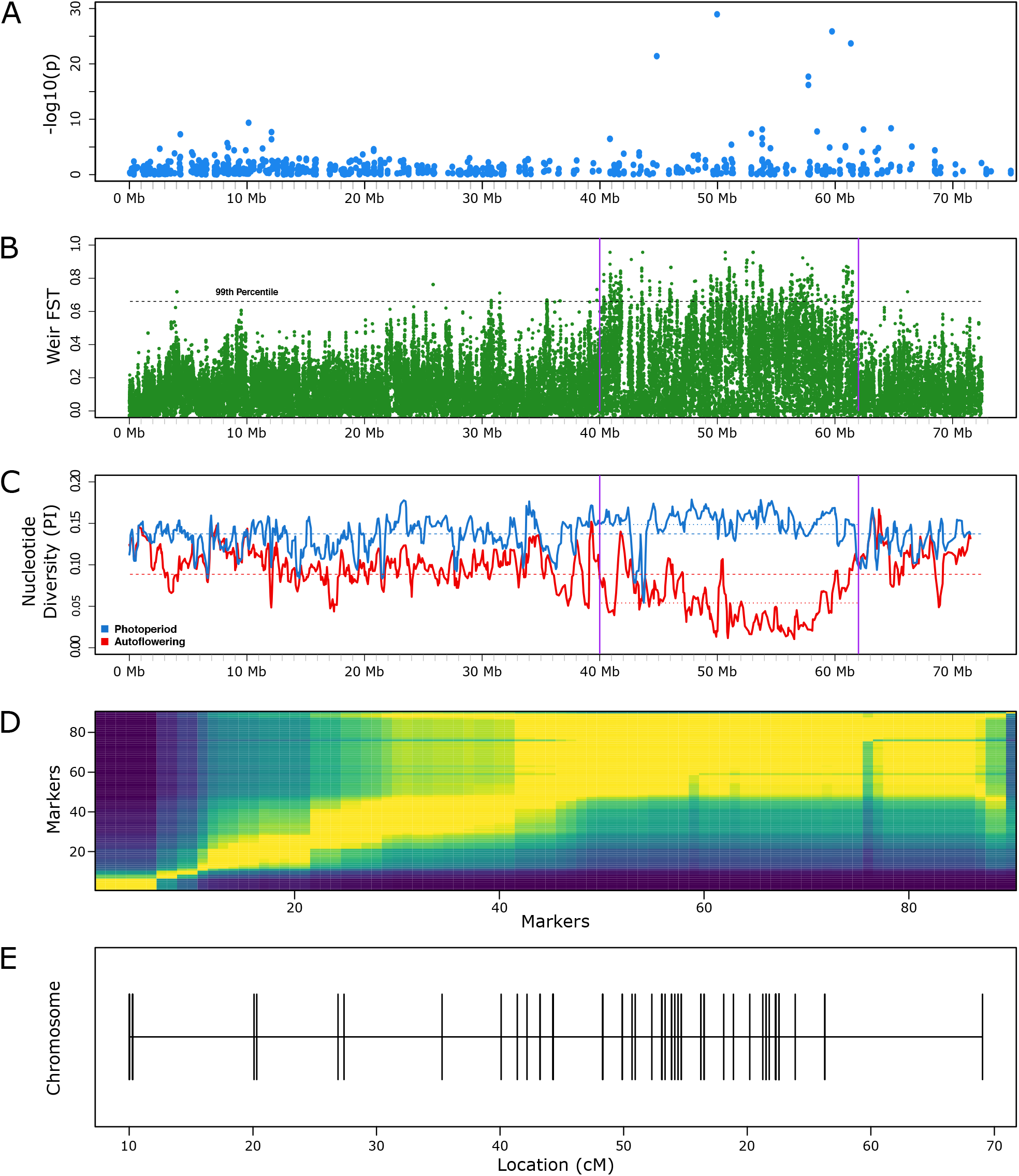
Mapping the auto-flowering trait to chromosome 5 in Cannabis. **(A)** Manhattan plot of 804 GBS SNPs, based on association mapping of 47 photoperiod-independent and 231 photoperiod cannabis accessions phenotypically encoded as 1 and 0 respectively. **(B-C)** Comparative population genetics statistics using 916K variants from WGS data of 12 autof-lowering and 12 photoperiod sensitive genotypes. The auto-flowering associated region of interest (40 to 62 Mb) is delineated by vertical purple lines. Statistics were calculated using a sliding window of 5Kb and a 1Kb step size. **(B)** Fixation index (F_st_) between populations. A value of 0 implies no observable difference in allele frequency in each window of variants between the two groups, while a value of 1 suggests that observed genetic variation is explained entirely by population structure and that each group is “fixed” for different observed alleles in that region. The 99th percentile of Fst values observed across all windows (F_ST_ = 0.66) is denoted with a dashed line. Of the 587 windows greater than or equal to the 99th percentile of all windows, 566 windows (96.4%) are found between 40 and 62 Mb. **(C)** Nucleotide diversity (Pi) comparing the average number of nucleotide differences between all possible pairs. Dashed lines represent the mean nucleotide diversity of all windows spanning the chromosome (Photoperiod = 0.137, Auto-flowering = 0.089), while the dotted lines represent the mean nucleotide diversity of windows located between 40 and 62 Mb: Photoperiod = 0.149 (increase of 8.76% relative to entire chromosome), Auto-flowering = 0.054 (decrease of 39.33% relative to entire chromosome). **(D)** Heat map of pairwise recombination fractions estimated with 90 markers in Chr 5 in an F2 population segregating for the auto-flowering trait. Low estimates of recombination fraction and high LOD scores are shaded in yellow, while blue represents the opposite. **(E)** Marker distances in cM calculated using pairwise recombination fraction and he Kosambi mapping function.

To gain more insight into the genetic diversity of the region identified by our GWAS study, whole genome sequencing (WGS) was conducted on 12 photoperiod sensitive and 12 auto-flowering genotypes. Using the SNPs observed between photoperiod sensitive and auto-flowering genotypes we calculated the fixation index (F_ST_) between the two populations. We found that the Fst values were significantly higher in a 22Mb region of chromosome 5 (Figure 1b). Nucleotide diversity (Pi) within this region was lower in the auto-flowering population compared to the photoperiod sensitive one (Figure 1c). To confirm that the observed differences in Fst and Pi values were true regardless of the reference genome, mapping was also conducted on the CS10 CBDRx genome assembly (Grassa et al., 2021). Similar genetic differences between the two populations were observed at the corresponding region within the CBDRx genome on chromosome NC_044371.1 (annotated as chromosome 1 in the CBRx reference assembly) (data not shown). For clarity, we will refer to chromosome 5 as the auto-flowering locus containing chromosome, in line with the Purple Kush annotation.

Our GWAS and population genetic approaches both identified a large chromosomal region associated with the auto-flowering trait, indicating either strong sequence divergence, and/or a region of low recombination. To resolve this, and to help find the causal variant, we constructed a genetic map of chromosome 5 with 192 individuals from an F2 population segregating for the auto-flowering locus. The observed recombination frequencies and genetic distances indicated the presence of a very low recombination region with the same coordinates as the region with significant F_st_ and Pi differences (Figure 1d-e).

### Fine Mapping the Auto-flowering Loci

The 22Mb region in chromosome 5 associated with the auto-flowering trait had 1353 genes representing a major challenge to identify the causal mutation. In order to narrow down the number of genes, we generated multiple F2 populations using diverse auto-flowering and photoperiod sensitive parents with the aim to find recombinants with a smaller region associated with the auto-flowering trait. A total of 7 F2 families were genotyped to identify rare recombination events in the low recombination region on chromosome 5. After screening 872 plants we found only 23 plants from two F2 populations that had recombination events in the 22MB region. The other 5 F2 families had no recombination events in the 22 MB region. With these results we were able to reduce the location for the proposed auto-flowering locus to a ~2.5Mb region located between 41,089,010 – 43,573,228bp (Purple Kush assembly).

From the F2 populations, four plants without the auto-flower phenotype were determined to be heterozygous for markers 1-3 and homozygous for markers 4-6, indicating a recombination event between markers 3 and 4. Plants were self-pollinated and two of the resulting populations were grown under long days to reveal that 4/24 and 7/44 plants exhibited the auto-flowering trait, conforming to the expected segregation ratio of 3:1. These results indicate that we were able to break the linkage region and refine the auto-flowering locus to the ~2.5Mb region between markers 1 and 4.

### Pseudo-response Regulator Differentially Expressed within Linkage Region

As fine mapping efforts were somewhat limited by the low recombination rates in this region, we used gene expression to identify possible candidate genes. The F2 population used for this experiment was segregating for the auto-flowering trait and importantly showed similar juvenile phase duration for auto-flowering and photoperiod sensitive plants. Furthermore, flowering time in auto-flowering genotypes did not significantly differ under LD compared to SD. Therefore, we focused on identifying gene candidates within the linkage region that are differentially expressed under LD vegetative growth. We hypothesized that the auto-flowering causal gene should have an expression pattern that meets the following two conditions: (1) is significantly different between auto-flowering and photoperiod sensitive genotypes in vegetative stage; and (2) show no differences in expression between auto-flowering and photoperiod sensitive during flowering.

RNA-Seq was used to measure gene expression on the above mentioned F2 population segregating for the auto-flowering phenotype. To validate the phenotype, a change from LD to SD was carried out once all auto-flowering carrying plants had terminal flowers. Out of 54 plants, 16 flowered under LD. Due to the differences in development time, sampling times were chosen to adequately cover the extent of each population’s vegetative and flowering stages. Differential gene expression analysis was focused on the vegetative growth stages. Across the genome, 2,826 genes were significantly differentially expressed between auto-flowering and photoperiod sensitive genotypes (FC > 1.5 and p-value < 0.05), indicating large regulatory changes during vegetative growth (Figure 2a). Of these, we identified 10 genes with homology to known *Arabidopsis* floral regulators including 3 genes with homology to *FT*. These results indicate that auto-flowering in cannabis may be due to the loss of *FT* repression under LD.

**Figure 2.**
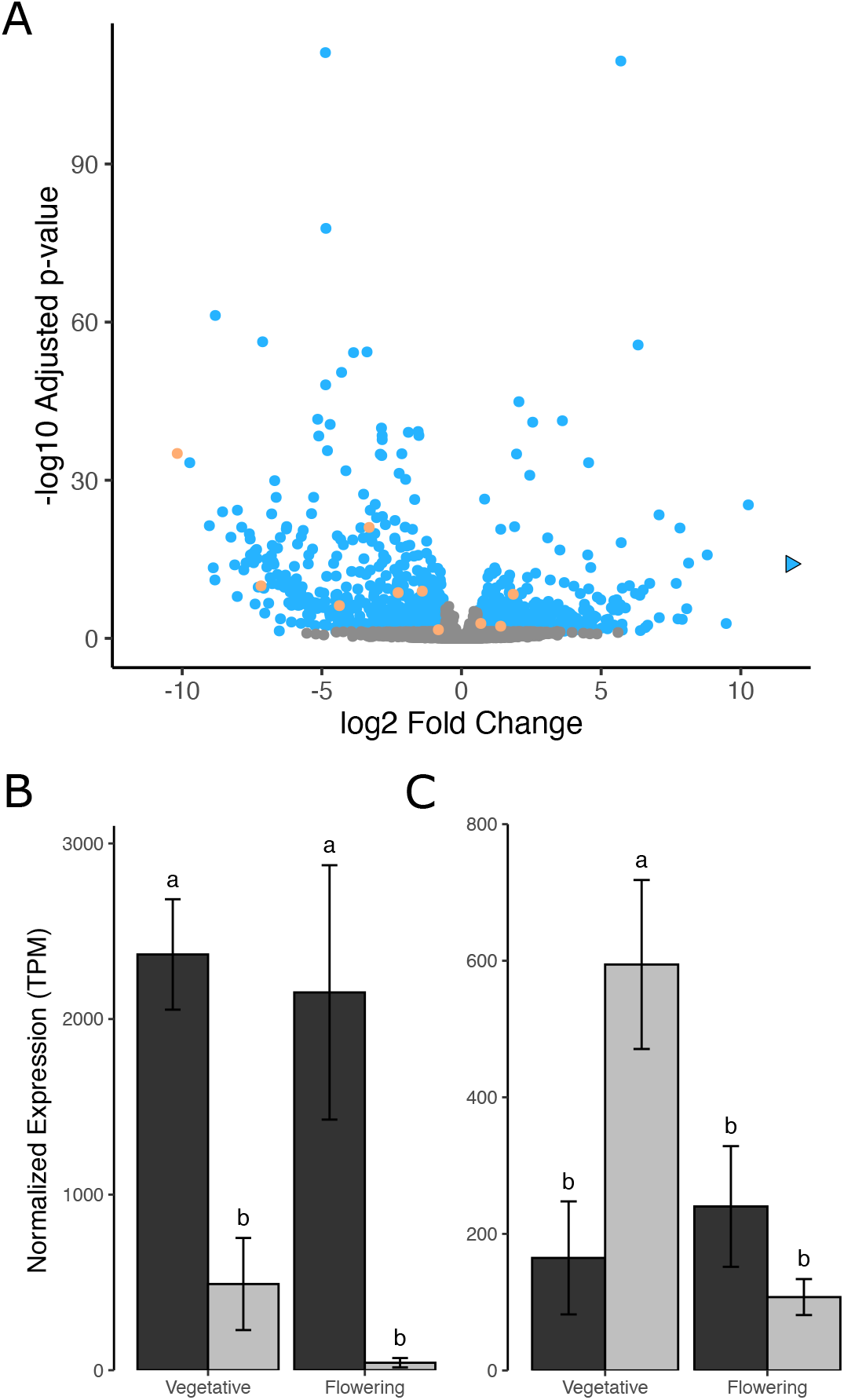
RNA-Seq differntial gene expression analysis. **(A)** Volcano plot depicting number of genes (N = 2,826) differentilly expression (DE) between photoperiod sensitive and auto-flowering genotypes during vegetative growth. Blue dots indicate signficantly DE genes (log2 fold change > 1.5, p < 0.05). Gray dots indicates genes not DE. Yellow dots indicate genes that are signficantly DE and have sequence homology to known *Arabidopsis* floral regulators. Blue arrow denotes point outside of plot at a log2 fold change of 23.5 **(B)** Mean normalized expression of *CsAP2* between auto-flowering (black) and photosensitive (gray) genotypes during vegetative and flowering stages. **(C)** Same as **(B)** but for *CsPRR37* expression.

Within the 2.5Mb region containing the auto-flowering locus, 2 genes were significantly differentially expressed and had homology to known *Arabidopsis* floral regulators. The first gene, LOC115708151, shows homology to Arabidopsis *APETALA 2 (AP2*), a floral homeotic gene involved in establishing floral meristem identity. The second gene, LOC115705128, is homologous to the *Arabidopsis* pseudo-response regulator family of circadian clock genes and is annotated as *PSUEDO-REPSONSE REGULATOR37(PRR37*). These genes will hereafter be referred to as *CsAP2* and *CsPRR37. CsAP2* expression was elevated in auto-flowering genotypes compared to photoperiod sensitive genotypes during vegetative growth (Figure 2b). During flowering, *CsAP2* remained elevated in auto-flowering plants while expression in photosensitive individuals was reduced to levels below those observed in vegetative growth. *CsPRR37* expression was markedly reduced in auto-flowering genotypes during vegetative growth compared to photosensitive genotypes (Figure 2c). During flowering, *CsPRR37* expression in photoperiod sensitive genotypes dropped to the same level observed in auto-flowering genotypes during vegetative growth. *CsPRR37* expression in auto-flowering genotypes did not significantly differ between vegetative and flowering stages.

It was unexpected to see *CsPRR37* expression in very young auto-flowering plants under “vegetative” conditions with low values which were very similar to photoperiod sensitive plants during flowering. This observation raised the question whether *CsPRR37* in auto-flowering plants could have a higher expression at a younger age before flowering is initiated. Or alternatively, *CsPRR37* could have low expression associated with flowering even at the seedling stage. To answer these questions, and gain further insight into both candidate genes, we performed an RT-qPCR experiment to profile the expression of flowering genes in leaves collected as early as possible (day 6 after germination). In addition to measuring expression of *CsPRR37*, we measured the expression of a highly induced *FT* homologue (LOC115697736) identified in the RNA-Seq analysis to see how early *FT* was being up regulated in auto-flowering plants. Interestingly, the selected *FT* homologue shows a greater homology to SD-specific rice florigen *HEADING DATE 3A (HD3A*), than any *Arabidopsis FT*.

### LD Expression of FT Homologue Observed in Auto-flowering Plants

Two populations, one auto-flowering and the other photoperiod sensitive, were grown from seed under LDs. Leaf samples were collected weekly until pistils were observed in auto-flowering genotypes on day 35, after which, plants were moved to SD with sampling increasing to twice a week. Elevated levels of Cannabis *Hd3a*, hereafter referred to as *CsHd3a*, were detected as early as 13 days post germination in auto-flowering genotypes. Expression levels increased rapidly between days 27 to 35 but fell back to pre-day 27 levels shortly after (Figure 3a). It should be highlighted here that the observed peak in auto-flowering *CsHd3a* expression occurred before moving plants into SD, with day 35 samples collected under LD. In comparison, photosensitive genotypes had nearly undetectable levels of *CsHd3a* mRNA until they were changed over to SD after which a more moderate spike in expression was observed (Figure 3b). These results support *CsHd3a* acting as a cannabis florigen and suggest a loss of repression of *CsHd3a* under LD in auto-flowering genotypes.

**Figure 3.**
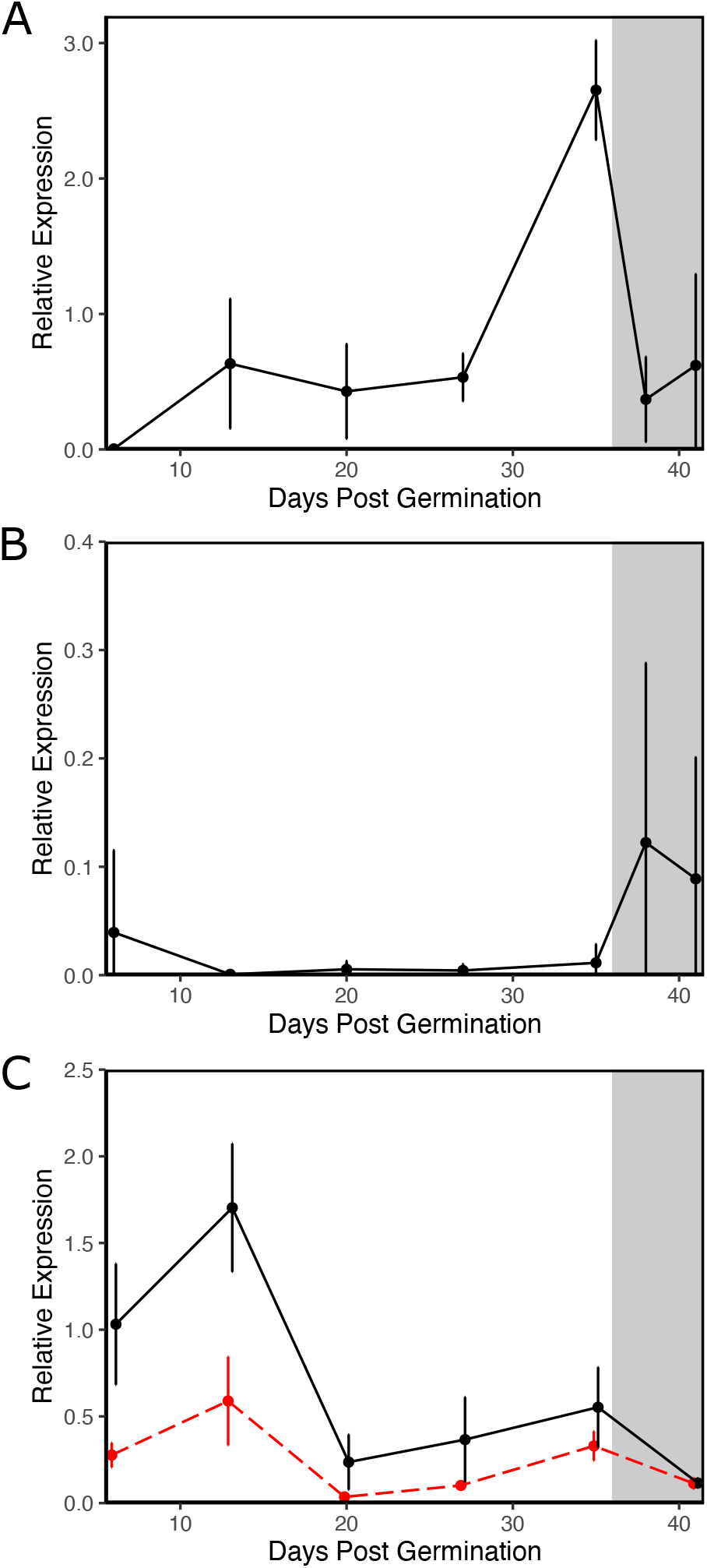
Mean expression levels of *CsHd3a* and *CsPRR37* mRNA throughout development. Gray areas represent days under SD (12h:12h), while white areas indicate days under LD (18h:6h). Biological replicates were a minimum of N = 3, maximum of N = 4. **(A)** *CsHd3a* expression in auto-flowering plants. **(B)** *CsHd3a* expression in photoperiod sensitive plants. **(C)** *CsPRR37* expression. Black line indicates photoperiod sensitive plants and red dashed line auto-flowering plants.

Auto-flowering *CsPRR37* mRNA levels remained well below photosensitive *CsPRR37* under LD conditions, particularly in the first two weeks of development (Figure 3c). After transitioning to SD, *CsPRR37* levels did not differ between auto-flowering and photoperiod sensitive plants. These results suggest that *CsPRR37* expression in leaves is altered during early “vegetative” growth but is expressed at levels similar to photoperiod sensitive plants during flowering, an observation consistent with our previously stated hypothesis. Since fine mapping efforts were unable to break the linkage between *CsPRR37* and *CsAP2*, we assessed its expression pattern throughout development. *CsAP2* mRNA levels steadily dropped over a five-week period of vegetative LD growth in photoperiod sensitive plants and continued to drop during SD (data not shown). In contrast, *CsAP2* expression in auto-flowering plants fluctuated during development and was only significantly different compared to photoperiod sensitive plants on days 6, 27, and 41.

### Mutation in CsPRR37 Abolishes Proper Splicing in Auto-flowering Genotypes

To further garner evidence for *CsPRR37* as the genetic source of the auto-flowering phenotype, we analyzed the WGS data from Figure 1 to identify structural variants associated with auto-flowering genotypes compared to photoperiod sensitive. No variants associated with the auto-flowering phenotype were identified in *CsAP2*. However, a G to T mutation was identified in *CsPRR37* that was present in all 12 auto-flowering genotypes sequenced and none of the 17 photoperiod genotypes. Further inspection showed that the mutation was present in the donor splice site of the second protein coding exon thereby abolishing the canonical GT donor splice site motif.

To confirm that the identified splice site mutation disrupts *CsPRR37* splicing in auto-flowering genotypes, full-length *CsPRR37* transcripts were amplified from cDNA from three different autoflowing genotypes, blunt ligated into a pJET vector and transformed into *Escherichia coli*. Multiple *E coli* colonies were selected for sanger sequencing of *CsPRR37*. Additionally, full-length *CsPRR37* transcripts from 2 photoperiod sensitive genotypes were also sequenced to verify canonical splicing. A single *CsPRR37* transcript variant was identified in photoperiod sensitive genotypes, matching NCBI’s annotation and our RNA-Seq results (Figure 4a). For auto-flowering genotypes, 5 transcript variants were identified, all different from the canonical variant present in photoperiod sensitive genotypes.

**Figure 4.**
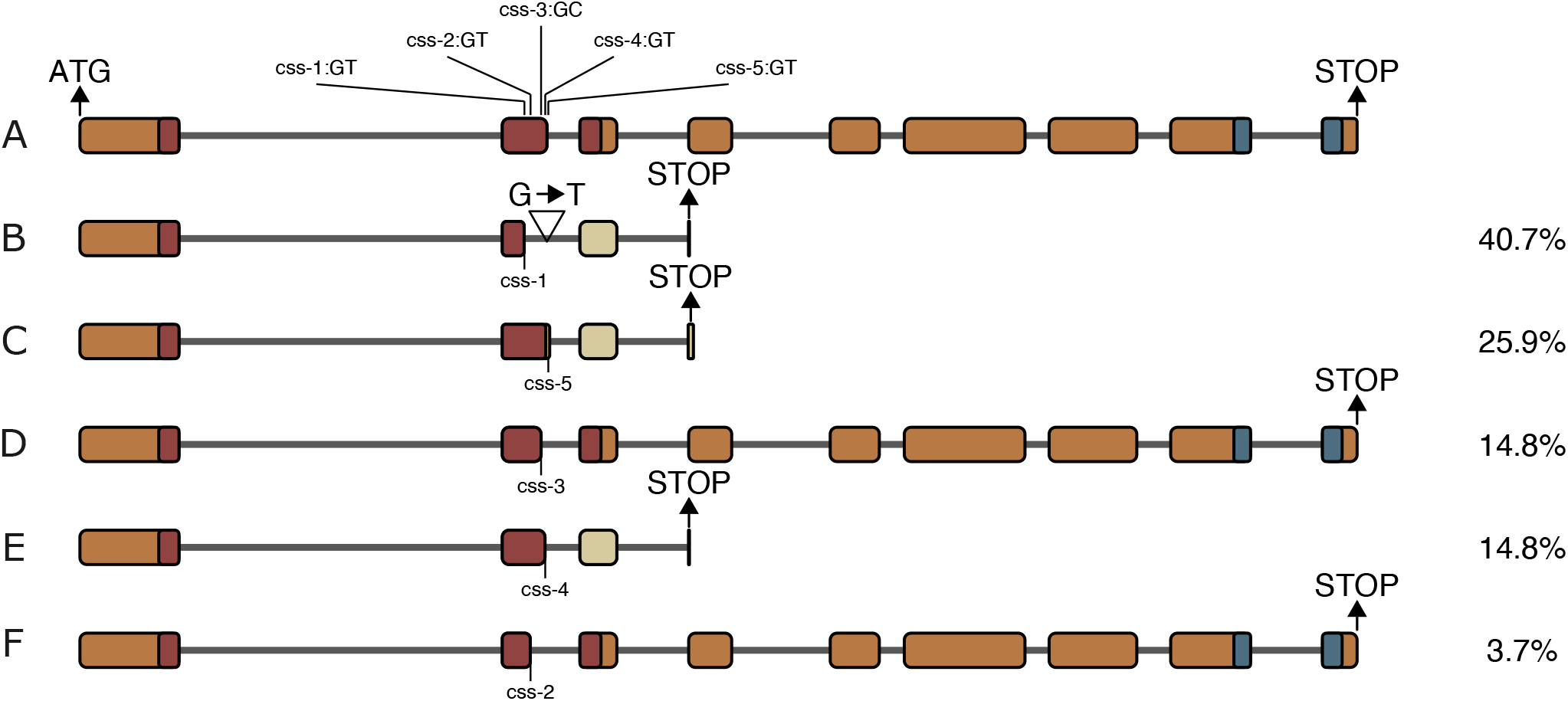
*CsPRR37* mRNA transcipt variants identified in photoperiod sensitive and auto-flowering plants. **(A)** Functional transcript identified in photoperiod sensitive plants. Position and sequence of cryptic splice sites used in transcripts isolated from auto-flowering plants are labeled on exon 2. **(B-F)** Transcript variants identified in auto-flowering plants. Position of G to T mutation knocking out the donor splice site after exon 2 is depicted in B and is present in all auto-flowering transcript variants. Variant abundence in auto-flowering plants is labeled on the right side of each transcript (N = 27). Introns are shown as solid lines and exons as boxes. Orange boxes indicate in frame exons and beige boxes as frame shifted exons. Red and blue boxes indicate positions of in frame PR domain and CCT domain respectively.

Auto-flowering transcript variants differed in their use of nearby cryptic splice sites, labeled css-1 to css-5. The most abundant variant, representing 40.7% of all transcripts sequenced, used css-1 located midway through the second exon at 1603bp from the translational start site (Figure 4b). The cDNA sequence from transcript variant 1 shows a 119bp deletion from the second exon producing an early stop codon due to a frameshift. The second most abundant transcript variant (25.9% of all variants sequenced) used css-5 located 1689bp from the TSS, 4bp past the end of the second exon (Figure 4c). The predicted protein contains a frameshift at the end of exon 2 due to the 4 bp insertion. A premature stop codon truncates the protein at the beginning of the 4^th^ exon. The third variant identified (14.8% of sequenced transcripts) produces a full-length protein with a deletion of 7 amino acids due to the use of a noncanonical GC cryptic splice site (css-3) located 1664bp from the TSS in the middle of exon 2 (Figure 4d). The fourth variant identified (also 14.8% of all sequenced transcripts) contains a 7bp deletion from the use of css-4 located 1678bp from the TSS within exon 2. The resulting protein contains a frame shift following the cryptic splice site junction and a premature stop codon at the start of exon 4 (Figure 4e). The last variant, present in only 3.7% of transcripts, produces a near full-length protein but contains a 20 amino acid deletion from the use of ccs-2, located 1625bp from the TSS in exon 2 (Figure 4f). All transcript variants identified in auto-flowering genotypes result in a protein with either a deletion or frame shift within the pseudo-response (PR) domain located within exons 2 and 3. The least abnormal transcripts (3 and 5), are likely translated into proteins with limited functionality due to the deletions in the PR domain. Transcripts 1, 2 and 4 are additionally missing their *CONSTANS, CONSTANS-LIKE*, and *TOC1* (CCT) domain due to premature stop codons at the beginning of the 4th exon. These results indicate that *CsPRR37* is likely non-functional in autof-lowering genotypes and strongly suggest it is the underlying gene responsible for the *Autof-lowering1* QTL described by Toth et al. (2022).

### LD clock in auto-flowering Cannabis is disrupted

We next investigated whether the splice mutation identified in *CsPRR7* could disrupt the circadian clock within auto-flowering cannabis plants thereby effecting flowering time. Pseudoresponse regulators are important components of the circadian clock and are characterized by an oscillating expression pattern peaking at specific times of the light-dark cycle. To determine if *CsPRR37* exhibits the expected oscillating expression pattern, RT-qPCR was used to measure *CsPRR37* gene expression every 3 hours over a 24h period. Sampling was conducted on both photoperiod sensitive and auto-flowering genotypes grown under LD (18h light, 6h dark). *CsPRR37* exhibited a bi-phasic expression with a midday peak at Zeitgeber Time (ZT) 11 and nighttime peak between ZT 20 and ZT 23 (Figure 5a). The nighttime peak was approximately twice that of the midday peak. The expression profiles between auto-flowering and photoperiod sensitive *CsPRR37* only differed in the amplitude of their midday peak where auto-flowering *CsPRR37* showed a 50% reduction in expression compared to photoperiod *CsPRR37*. We hypothesized that in addition to the midday peak differences in *CsPRR37* transcript levels, its predicted truncated protein products should disrupt the expression of other clock components, in particular, those directly regulated by *CsPRR37*.

**Figure 5.**
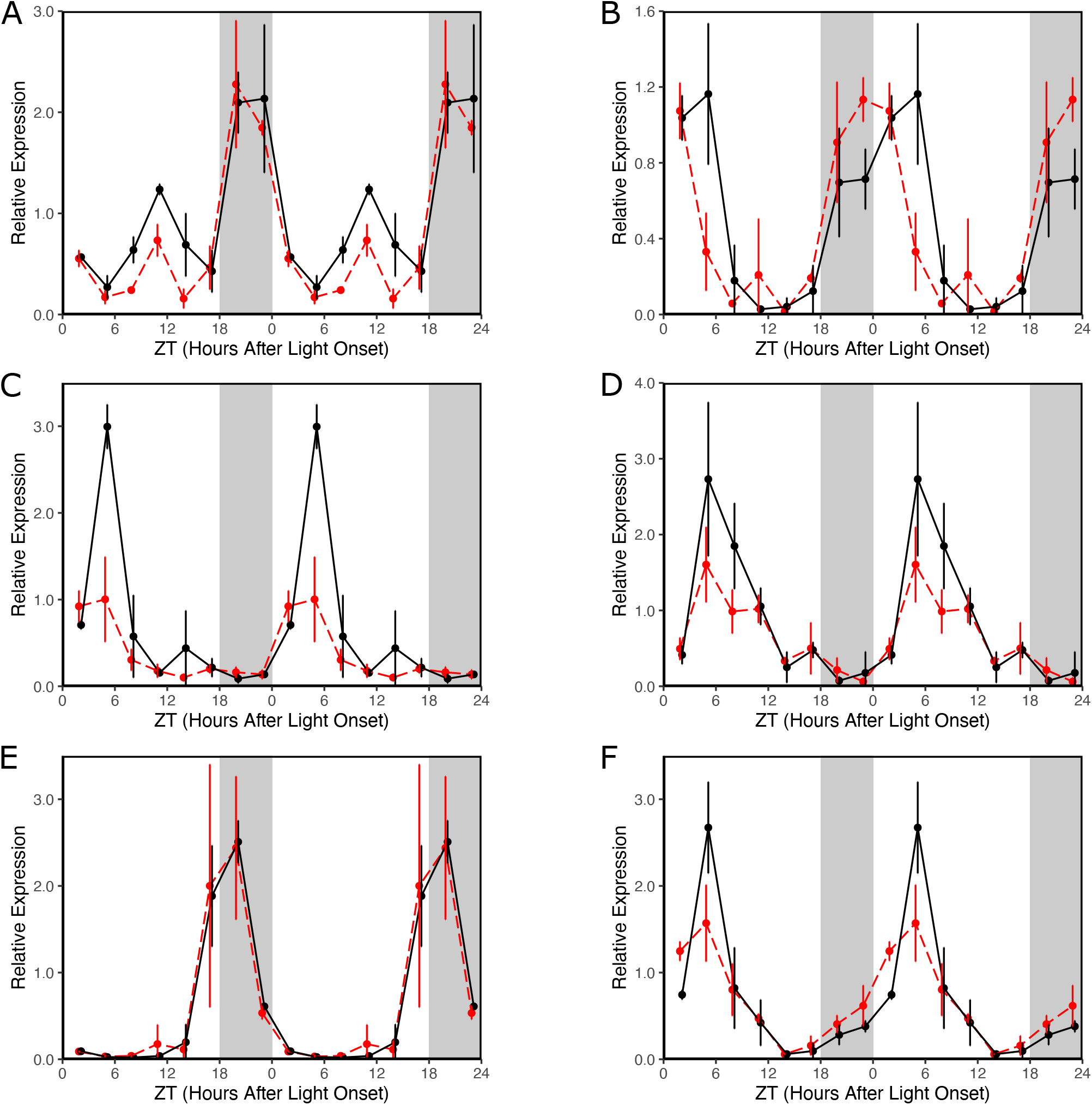
Mean mRNA expression of circadian clock genes **(A)** *CsPRR37*, **(B)** *CsPRR95*, **(C)** *CsPRR3*, **(D)** *CsTOC1/PRR1*, **(E)** *CsLHY*, and **(F)** *CsGI* over a 24h period under LD (18:6h) photoperiod. White and gray areas represent white light and dark conditions respectively. Data is repeated once to better visualize oscillation patterns. Black lines indicate photoperiod sensitive plants and red dashed lines indicate auto-flowering plants. A minimum of N = 3 and maximum of N = 4 biological replicates were used. An artifical nudge was applied to x-axis coordinates to distinguish between overlapping observations.

To explore the potential effects in the clock of non-functional and truncated auto-flowering CsPRR37 proteins, we measured expression of other PRR genes in the cannabis clock, namely *CsPRR95* (LOC115705125), *CsPRR3*(LOC115716206), and *CsPRR1*(LOC115711591) over a 24-hour period entrained under LD. Cannabis *LATE ELONGATED HYPOCOTY (CsLHY*, LOC115706434), a Myb-related gene with homology to *Arabidopsis* morning complex gene, and circadian regulated gene *GIGANTEA (CsGI*, LOC115708742) were included. *CsPRR95* exhibited a monophasic expression pattern peaking at ZT 5 in photoperiod sensitive genotypes (Figure 5b). A 6-hour phase advance was observed for *CsPRR95* in auto-flowering genotypes compared to photoperiod sensitive with peak expression at ZT 23. *CsPRR3* expression was monophasic, peaking at ZT 5 in both photoperiod sensitive and auto-flowering genotypes, however, its peak expression was significantly lower in auto-flowering genotypes compared to photoperiod sensitive (Figure 5c). Additionally, auto-flowering *CsPRR3* expression appeared to increase and decrease slightly earlier compared to the photoperiod sensitive, suggesting a possible phase advance. *CsPRR1* oscillation was monophasic, with a sharp increase in expression peaking at ZT 5 followed by a gradual decrease, characteristic of *PRR1/TOC1* expression observed in other plant species (Figure 5d). Peak expression in *CsPRR1* appeared to be dampened in auto-flowering plants compared to photoperiod sensitive, however this was not statistically significant. *CsLHY* expression was again monophasic, with peak expression at ZT 20 (Figure 5e). No differences were observed between auto-flowering and photoperiod sensitive genotypes for *CsLHY. CsGI* expression peaked at ZT 5 in both auto-flowering and photoperiod sensitive genotypes (Figure 5f). Peak expression in auto-flowering *CsGI* plants was significantly dampened compared to photoperiod plants, with auto-flowering mRNA accumulating earlier in the day, suggesting a possible phase advance. These results indicate that in total four core components the circadian clock in auto-flowering cannabis, CsPRR37, CsPRR3, PRR95, and *CsGI* are significantly disrupted under LD. Importantly, *CsPRR37* is the only gene which showed impairing mutations in auto-flowering genotypes whereas other clock genes investigated here had identical predicted protein sequences between auto-flowering and photoperiod sensitive genotypes.

### SD Photoperiods Exhibit Consecutive Waves of PRR Gene Expression

We next investigated whether the cannabis clock genes exhibit altered oscillation patterns under SD conditions (12 hours light/12 hours dark) compared to LD, and if oscillation patterns would differ between auto-flowering and photoperiod sensitive plants. Plants entrained under SD were sampled over a 24-hour period and mRNA expression of circadian clock genes was measured. In photoperiod sensitive plants, *CsLHY* expression peaked between ZT −1 and ZT 2, likely reaching its maximum around ZT 0 (Figure 6a). *CsPRR37* was upregulated between ZT −1 and ZT 5. It should be noted that at ZT 2, two of the four biological replicates peaked in expression while the remaining 2 were depressed, indicated by the large standard deviation associated with mean expression at ZT 2. *CsPRR95, CsPRR3*, and *CsPRR1/TOC1* expression peak at ZT 5, ZT 8, and ZT 11 respectively. In auto-flowering plants, similar expression patterns were observed with a few notable differences. *CsPRR37* peak expression was more apparent at ZT 2, and *CsPRR1/TOC1* expression had a 3-hour phase advance, peaking at ZT 8 (Figure 6b). Moreover, *CsPRR95* and *CsPRR3* expression peaks were not as sharp as in photoperiod sensitive plants, possibly indicating a phase advance not captured by the sampled timepoints. These result show that clock gene oscillations are dramatically different in SD photoperiods compared to LD. Furthermore, circadian clock gene regulation under SD differs between photoperiod sensitive and auto-flowering plants, albeit not as pronounced as under LD.

**Figure 6.**
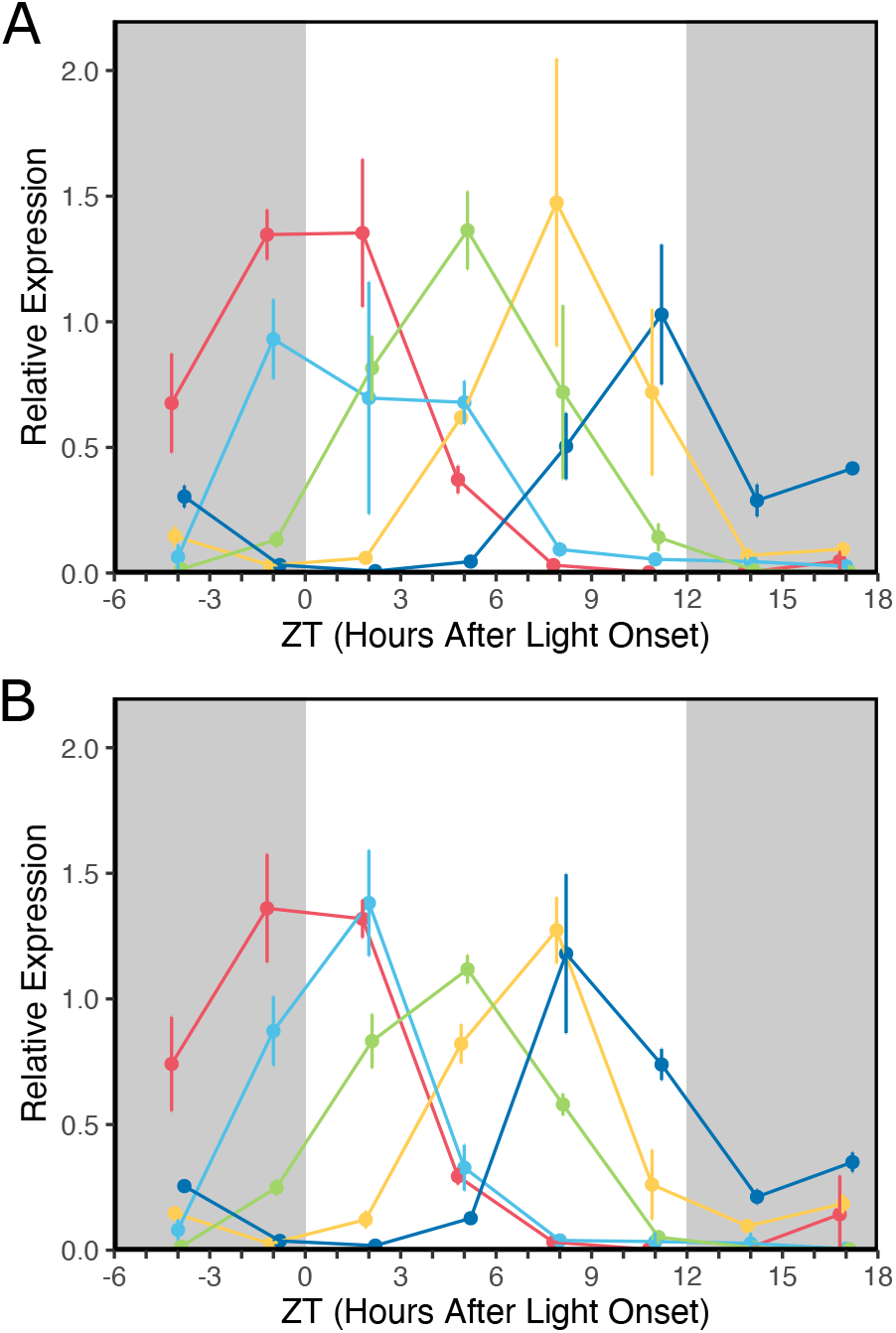
Mean mRNA expression patterns of circaidan clock genes over a 24 hour period under SD photoperiod (12h light/12h dark). **(A)** Photoperiod sensitive plants and **(B)** auto-flowering plants. Circadian clock genes in order of expression peaks are *CsLHY* (red), *CsPRR37* (light blue), *CsPRR95* (green), *CsPRR3* (yellow), and *CsTOC1/PRR1* (dark blue). White and gray bars represent white light and dark conditions respectively. A minimum of N = 3 and maximum of N = 4 biological replicates were used. Vertical lines indicates standard deviation.

## Discussion

### Circadian Clock Genes Drive Domestication

Manipulating flowering time is essential to grow a crop at novel latitudes, therefore, understanding the specific functions of genes that control photoperiod sensitivity is critical for domestication of new crops and efforts to expand the cultivation range of existing crops. Genes involved in, or closely related to the circadian clock, are well conserved in function across many plant species (Searle and Coupland, 2004). Pseudo-response regulators (PRR) are well conserved circadian clock genes commonly identified in flowering time QTLs of domesticated crops including Barley, Wheat, Sorghum, Rice, and Beets. The Ppd-H1 allele in barley encodes a pseudo-response regulator homologous to Arabidopsis *PRR7*. The recessive ppd-H1 allele results in late heading allowing spring sown varieties of barely to increase yields (Turner et al., 2005). The closely related wheat ppd-D1a allele results in photoperiod insensitivity by misexpression of a PRR resulting in the up regulation of the wheat homologue of *FT*, *TaFT1* (Beales et al., 2007). Similarly, in SD heading crop Sorghum, multiple independent mutations in *SbPRR37* are responsible for the early heading *ma1* allele increasing *SbFT* expression under LD (Murphy et al., 2011). In rice, heading date major effect QTL *EH7-2/Hd2* was identified as *OsPRR37* (Murakami et al., 2005; Koo et al., 2013). Our experiments add to this list of key PRR domestication mutations, with GWAS, population genetic analysis, fine mapping and gene expression data all indicating the involvement of a mutated CsPRR37 allele in the loss of photoperiod sensitivity in cannabis.

### Adaptation to Higher Latitudes may be Attributed to CsPRR37 Variation

Photoperiod insensitivity is a common trait observed in SD crops domesticated in higher latitudes. Intensive artificial selection in rice has allowed its adaptation to latitudes as high as 53°N. Rice grown at these latitudes flower extremely early with weak or no photoperiod sensitivity (Wei et al., 2008; Fujino and Sekiguchi 2005). Crosses between late flowering *indica* cultivars with early flowering *japonica* have identified many distinct heading date QTLs contributing to a diverse range of flowering times (Yoo et al., 2007; Uga et al., 2007; Doi et al., 2004; Monna et al., 2002). A major effect QTL *Hd2* was identified as *OsPRR37* (Murakami et al., 2005; Koo et al., 2013). Koo et al. (2013) showed that natural variation in *OsPRR37* contributed to the expansion of rice cultivation in northern latitudes, where cultivars grown at the most northern regions contained loss-of-function alleles.

Recently, Chen et al., (2022) collected both natural and domesticated accession of cannabis across China. Flowering time experiments showed that wild collections flowered under LD. All natural accessions except for two, were collected in northern China at latitudes ranging from 41.50 – 50.16°N. The two which were not from northern China were collected in northern Yunnan and Xizang province, close to the Tibetan plateau where cannabis is proposed to have originated (McPartland et al., 2019). Gene expression analysis of florigen *CsHd3a* showed strong upregulation under LD in wild accessions, similar to our observations in Figure 3a. Furthermore, an analysis of nucleotide diversity between domesticated photoperiod sensitive and wild accessions identified genes under positive selection, of which *CsPRR37* was significant. Like rice, natural variation in *CsPRR37* may be involved in cannabis adaptation to varying latitudes by regulating flowering time, with loss-of-function alleles present in the highest latitude regions. A detailed analysis of *CsPRR37* in wild and domesticate genotypes adapted to varying latitudes will be required to confirm this hypothesis. Currently, sequence data from auto-flowering wild accession collected by Chen et al., (2022) are under embargo, however, wild accession collected by Ren et al., (2021) overlap geographical regions. While flowering time was not analyzed in this study, the search for *CsPRR37* mutation(s) within these accessions is underway.

### CsPRR37 may repress CsHd3a, preventing flowering under LD

Photoperiod insensitive genotypes of cannabis were found to contain a splice site mutation in *CsPRR37. CsPRR37* is homologous to other PRR family genes within the PRR3/7 clade (Hotta et al., 2022), many of which have been confirmed as *bona fide* timekeepers such as *PRR7* in *Arabidopsis*, and *PRR37* in rice. *CsPRR37* contains both PR and CCT domains which are signature features of pseudo-response regulators. The CCT domain is involved in nuclear localization and DNA binding (Tiwari et al., 2010; Robson et al., 2001). The PR domain is involved in forming heterodimers (Yuan et al., 2021). Sequencing of *CsPRR37* transcripts from auto-flowing genotypes identified 5 variants, 3 of which had early stop codons and lacked a CCT domain. The other two transcript variants had deletions in the PR domain and an apparently functional CCT domain. These results suggest that the *CsPRR37* splice site mutation identified in auto-flowering plants likely causes a severe loss of function in the CsPRR37 protein due to a combination of lower gene expression and disrupted PR and CCT domains.

In *Arabidopsis, PRR7* forms a regulatory feedback loop repressing *CCA1* and *LHY* expression in conjunction with *PRR9* and *PRR5* (Nakamichi et al., 2010; Farre et al., 2005). Double and triple PRR mutants involving non-functional PRR7 has been shown to disrupt *CCA1* and *LHY* circadian rhythm effecting flowering time (Nakamichi et al., 2007). Under both LD and SD conditions no differences in were observed in *CsLHY* regulation between auto-flowering and photoperiod sensitive genotypes. This may be explained by the semi-redundant function of PRR genes, as observed in *Arabidopsis*, but doesn’t explain the early flowering phenotype of auto-flowering cannabis. Alternatively, *CsPRR37* may have a more direct function in regulating flowering as it does in other SD flowering crops. In rice, OsPRR37 delays heading date by repressing *Hd3a* under LDs independent of key photoperiod regulators *Hd1* (a CONSTANS ortholog) and *Ehd1* (Koo et al., 2013; Yano et al., 2004). A similar regulation in cannabis would explain auto-flowering where a non-functional CsPRR37 would fail to repress *CsHd3a* under LD. Indeed, *CsHd3a* expression is upregulated in auto-flowering plants in LD. It is interesting to note that the *FT* homologue we analyzed has a closer homology to rice *Hd3a*, which is a SD specific florigen as opposed to rice *RFT1*, a LD specific florigen.

### Circadian Rhythms are Drastically Altered between SD and LD Photoperiods

Cannabis circadian clock genes were sequentially expressed during SD photoperiod between ZT 0 (lights on) and ZT 12 (lights off). Peak expression of clock genes occurred approximately every 3 hours in the following order: *CsLHY*, *CsPRR37*, *CsPRR95*, *CsPRR3*, and *CsPRR1/TOC1*. This same pattern has been observed in the circadian clock genes of both *Arabidopsis* and Rice, highlighting its conservation in higher plants (Murakami et al., 2003; Matsushika et al., 2000). Unexpectedly, this pattern deviated drastically when cannabis was grown under LD (18h light and 6h dark). Under this photoperiod *CsTOC1*, *CsPRR95*, and *CsPRR3* all have overlapping oscillations with peak expression at ZT 5. Most striking was the inverted oscillation peaks observed in *CsLHY* and CsPRR1/TOC1. In *Arabidopsis*, the regulatory feedback loop created by morning expressed gene *LHY* (and homologous *CCA1*) and evening expressed *TOC1* composes the core oscillator of the circadian clock (Alabadi et al., 2001). This feedback loop appears to be conserved in other plant species (Wang et al., 2011; Murakami et al., 2007). In LD entrained cannabis, *CsLHY* peak oscillation occurred at night (ZT 20), while *CsTOC/PRR1* expression peaked in the subjective morning at ZT 5. It should be noted that an 18h:6h photoperiod is highly unnatural for cannabis and was chosen because of its use in the commercial production of drug-type cannabis. Therefore, the atypical oscillations observed under 18h:6h photoperiod may be the results of the circadian clock operating under a longer daylength than it evolved to do so.

### CsPRR37 Biphasic Expression may be Light Dependent

Except for the phase advance observed in auto-flowering CsPRR1/TOC1, circadian rhythms between auto-flowering and photoperiod sensitive plants did not drastically differ under SD. During LD, both auto-flowering *CsPRR95*, *CsPRR3*, and *CsGI* oscillations were altered. While not statistically significant, *CsPRR1/TOC1* peak expression was attenuated in auto-flowering plants under LD as well. The increased perturbation of circadian clock genes in auto-flowering plants in LD compared to SD may allude to a LD specific function of *CsPRR37*. Interestingly, the daytime peak of *CsPRR37* seemed to fill a gap in the circadian clock, being the only gene upregulated between ZT 9 and ZT 14. Moreso, *CsPRR37* was the only circadian clock gene analyzed in this study that showed a biphasic expression pattern observed only under LD. *CsPRR37* expression showed peaks at ZT 11 (daytime) and ZT 20 - 23 (nighttime). A biphasic expression pattern has been reported in Sorghum, with *SbPRR37* similarly being expressed at ZT 3 and ZT 15 under LD photoperiods (Murphy et al., 2011). Under SD, *SbPRR37* loses its evening peak. Constant light (LL) experiments to assess the free-running period of *SbPRR37* found that *SbPRR37* expression is light dependent, with both morning and evening peaks present under LD and LL, while SD and constant dark (DD) conditions yielding only the morning peak. Murphy et al., (2011) demonstrated that *SbPRR37* is a central repressor in the flowering regulatory pathway in response to photoperiod, with both morning and evening peaks needed to sufficiently repress expression of *SbFT*. Given the similarities in *CsPRR37* expression, LL and DD experiments are currently being conducted to assess the light dependency of *CsPRR37*. These experiments will additionally be used to better understand the free running periods of circadian clock genes analyzed in this study, and how integral *CsPRR37* is to the clock’s proper function.

## Conclusion

The large introgression carried in our diverse panel of auto-flowering genotypes (Figure 1) is likely shared with most auto-flower accessions in cannabis and explained by the severely reduced recombination observed between photoperiod sensitive and auto-flowering haplotypes. Currently available auto-flowering genotypes are known to have lower cannabinoid concentrations and poor flower quality than photoperiod sensitive cannabis, likely due to other QTLs on this large introgression. Knowing the causal mutation, which we believe is in *CsPRR37*, enables large scale screens for rare recombinants, as we have demonstrated, which we expect will break this linkage drag and facilitate the breeding of auto-flowering cultivars with competitive quality to photoperiod sensitive ones. Advancements in auto-flowering genetics will help transition the drug-type cannabis industry from clonally propagated indoor grown production to outdoor seed grown cultivars adapted to a diverse range of bioregions.

## Methods and Materials

The experiments outlined in this study were conducted using the Purple Kush reference genome. This includes all population genetics studies (unless stated otherwise), RNA-Seq, and primer design. However, due to the superior gene annotation provided on NCBI for the CBDRx genome, we relied on this reference for the functional annotation of the genes studied within this report.

### Plant Material and Growing Conditions

All plants used for gene expression assays (RT-qPCR and RNA-Seq) were grown from seed under artificial light. Seeds were sterilized in a 5% solution of PPM (Plant Preserve Mixture, Caisson Labs) for three hours, then transferred to petri dishes lined with Whatman filter paper saturated with 0.5% PPM. Seeds were left in the dark for approximately two days to allow for germination. After which, germinated seeds were transplanted into soil. SD photoperiod conditions consisted of 12 hours light and 12 hours darkness. LD photoperiod consisted of 18 hours light and 6 hours darkness.

Plants used in fine mapping the auto-flowering locus were grown hydroponically in a greenhouse using a mix of natural and artificial light. Seeds were germinated in rockwool, then transplanted into 4” rockwool cubes after two weeks. To produce F3 and F4 seeds, plants were self-pollinated by applying a foliar spray of 0.02 molar silver thiosulfate solution to induce male flowers to form on genetically female plants, then securing the plant in a pollination bag to prevent cross-pollination.

### Genome Wide Association Study

Genomic DNA libraries were prepared from purified DNA samples based on the methodology published in Gilchrist et al. (2021), based on Poland et al., (2012). Sequencing and variant calling was performed according to Gilchrist et al., (2021), using Trimmomatic (Bolger et al., 2014) for read trimming and the GBS-SNP-CROP 2.0 pipeline (Melo et al., 2016) for parsing, demultiplexing, alignment, and variant calling on the ASM23057v2 Purple Kush genome assembly (Laverty et al., 2019).

A Genome-Wide Association Study (GWAS) was performed using a Mixed Linear Model in TASSEL5 with 278 samples (47 auto-flowering, 231 photoperiod). The kinship (K) matrix was generated using centered Identity-By-State (IBS). The first 5 principal components of the genotype data were included in the model to control population structure. For mapping analysis, phenotype data was coded as 1 for auto-flowering and 0 for photoperiod sensitive.

### Genetic Diversity and Population Structure

Comparative population genetics statistics for chromosome 5 (Purple Kush assembly ASM23057v4) were performed using 916K variants from whole-genome sequencing (WGS) data from 17 photoperiod sensitive and 12 auto-flowering genotypes. Analyses were performed using a sliding window of 5,000 base pairs and variants contained within, with a step size of 1,000 base pairs across the chromosome. Fixation index (F_ST_) values (Weir & Cockerham, 1984) were calculated with VCFtools v0.1.16 (Danecek et al., 2011) to compare allele frequency differences (e.g., potential localized population structure) between 12 photoperiod-independent and 12 photoperiod accessions to have a balanced analysis. A value of 0 implies no observable difference in allele frequency in each window of variants between the two groups, while a value of 1 suggests that observed genetic variation is explained entirely by population structure and that each group is “fixed” for different observed alleles in that region. Nucleotide diversity (Pi) was calculated comparing the average number of nucleotide differences between all possible pairs within the 12 auto-flowering genotypes and the 12 photoperiod genotypes using TASSEL5.

### Genetic Map Construction

An F2 population was developed between and auto-flowering and photoperiod sensitive genotypes to construct a genetic map. A total of 192 F2 plants were grown indoor and genotyped using a SeqSNP chip with 5,000 custom markers selected from our GBS dataset to cover the entire cannabis genome. Biosearch Technologies performed the mapping of reads and SNP calling using the Purple Kush genome (ASM23057v4) as reference. In brief, variant calling was done using GATK and identified SNPs were filtered by biallelic variants. Only those SNPs for which a definitive parental origin could be assigned were used for genetic map construction. Of the 4004 markers available, only 484 remained after filtering for missing genotypes and segregation distortion. A linkage map was constructed with the R package Rqtl using the ‘est.map’ function with a genotyping error rate set to 0.01. Genetic distances were calculated from the observed recombination fractions using the Kosambi function.

### Fine Mapping of Auto-flowering Locus

Fine-mapping was done using Kompetitive Allele-Specific PCR (KASP) genotyping technology. All DNA samples for KASP genotyping were extracted from young leaf tissue on an Oktopure DNA extraction system using sbeadex Plant DNA purification kits (Biosearch Technologies). KASP assays were designed and produced by Biosearch Technologies based on provided sequence data for six locations distributed across the region of low recombination on chromosome 5. Genotyping assays were run on a QuantStudio (ThermoFisher) using low ROX KASP genotyping master mix (Biosearch Technologies).

Seven auto-flowering x photoperiod sensitive F2 populations were screened with the marker panel and only two showed individuals with recombination events in the 22 Mb region in chromosome 5. Recombination events were identified as a switch from heterozygous to homozygous genotype occurring between markers 1 and 6. The two Auto x Photo F2 populations that showed evidence of recombination were selfed to produce F3 progeny. Two plants heterozygous for the auto-flowering allele with a recombination event occurring between Markers 3 and 4 were self-pollinated to create segregating validation populations. The two recombinant populations (n=116) were grown under long day conditions and phenotyped for flowering time.

### RNA-Seq

RNA-Seq was performed on a F2 population segregating for the auto-flowering trait. Initially, leaf tissue from 54 plants grown under LD was collected every 2 to 3 days starting 17 days post sowing of seeds. Pistils were observed in plants carrying the auto-flowering on day 30. On day 46 of the experiment, plants carrying the auto-flowering trait were well into flowering and the remaining photoperiod sensitive plants were moved to SD. Sampling continued for photoperiod sensitive plants until day 56 when flowering was well into development. Two flowering and 2 vegetative timepoints were selected to perform RNA-Seq in auto-flowering plants. Two flowering and 3 vegetative timepoints were selected for photoperiod sensitive plants, with the extra vegetative timepoint selected to better cover the longer vegetative phase in photoperiod sensitive plants. Three biological replicates were included for both phenotypes.

Collected leaf tissue was flash frozen in liquid nitrogen immediately after sampling. Total RNA was extracted using an RNeasy Kit (Qiagen) and used for library preparation using Illumina’s TruSeq stranded mRNA kit. RNA sequencing was performed using Illumina’s NovaSeq6000 platform using 150-cycles paired-end sequencing. Sequence reads were quality checked using FastQC (Andrews, 2010) and trimmed to a minimum phred score of Q30 using Skewer (Jiang et al., 2014). Sequence reads were mapped to the Purple Kush reference assembly, NCBI accession ASM23057v4, using the HISAT2 v2.1.0 splice aligner (Kim et al., 2019). Mapped reads were assembled into transcripts and quantified for gene expression using Stringtie v2.1.4 (Kovaka et al., 2019). Differential gene expression was carried out using the DESeq2 R package v.1.20.1 (Love et al., 2014).

### Transcript Isolation and Sequencing

Three independent leaf samples were taken from auto-flowering genotypes and two from photoperiod genotypes for the purpose of RNA extraction. Plants were grown from seed and samples were taken at independent timepoints. Samples were collected in liquid nitrogen and placed at −80°C prior to RNA extraction. Total RNA was extracted using an RNeasy Plant Mini Kit from Qiagen. cDNA was synthesized using QuantiTect Reverse Transcriptase Kit from Qiagen.

PCR Amplification of the *CsPRR37* transcripts was done from predicted start codon to predicted stop codon, based on annotations of the Purple Kush genome. Additional amplification and sequencing was done to confirm the sequence of the C-terminal. PCR amplicons were ligated into a vector using the CloneJET PCR cloning kit and were transformed into E. cloni chemically competent *E. coli*. Plasmids were extracted from individual colonies using a QIAprep Spin Miniprep Kit from Qiagen. Plasmids were individually sanger sequenced using both CloneJET specific primers and primers internal to *CsPRR37*. Sequences showing multiple traces, or which were of low quality were not included in the analysis. A total of 27 independent high quality auto-flowering *CsPRR37* sequences were used to derive frequency rates of transcript variants.

### RT-qPCR

For the developmental time course study, leaf sampling was conducted approximately once a week for 6 weeks starting 6 days post seed germination. Sampling times were increased to twice a week after plants were moved to SD to better capture developmental changes. All samples were collected at the same time of the day.

For the 24-hour time course experiments, transplanted seeds were split into LD and SD photoperiods. Sampling of LD plants occurred 19 days after germination when both photoperiod and auto-flowering plants were in vegetative growth. For the SD condition, plants were sampled 61 days after germination during flowering. Under both conditions, leaf samples were collected every 3 hours for a 24-hour period.

All leaf samples were flash frozen in liquid nitrogen immediately after sampling. Total RNA was extracted using a RNeasy Plant Mini Kit from Qiagen. cDNA was synthesized using QuantiTect Reverse Transcriptase Kit from Qiagen. RT-qPCR was carried out using SsoAdvanced Universal SYBR Green Supermix from Bio-Rad on a CFX96 Deep-Well Real-Time system. A combination of two reference genes were used to normalize data. Initially, 5 genes were identified in the RNA-Seq study that had minimal variation between developmental timepoints and biological replicates. Reference genes stability across all developmental timepoints were evaluated using the GeNorm algorithm in Bio-Rad’s CFX Maestro Software (Vandesompele et al. 2002). All reference gene candidates were rated ideal with two genes, CsCUL3A (LOC115698686) and CsAGC (LOC115723274), being the most stable and used for this study. Evaluations on 24-hour timepoints were also conducted with similar stability results. All primers were designed based off of the RNA-Seq transcript positions when mapped to the Purple Kush reference genome. Primers were designed to capture all transcript variants determined by RNA-Seq, with one primer in each pair bridging an exon-exon boundary.

## Notes

### Competing Interest Statement

The authors declare financial competing interests by being employees of Aurora Cannabis Inc.

